# Taxonomy annotation and guide tree errors in 16S rRNA databases

**DOI:** 10.1101/288654

**Authors:** Robert C. Edgar

## Abstract

Sequencing of the 16S ribosomal RNA (rRNA) gene is widely used to survey microbial communities. Specialized 16S rRNA databases have been developed to support this approach including Greengenes, RDP and SILVA. Most taxonomy annotations in these databases are predictions from sequence rather than authoritative assignments based on studies of type strains or isolates. In this work, I investigated the taxonomy annotations and guide trees provided by these databases. Using a blinded test, I estimated that the annotation error rate of the RDP database is ~10%. The branching orders of the Greengenes and SILVA guide trees were found to disagree at comparable rates with each other and with taxonomy annotations according to the training set (authoritative reference) provided by RDP, indicating that the trees have comparable quality. I found 249,490 identical sequences with conflicting annotations in SILVA v128 and Greengenes v13.5 at ranks up to phylum (7,804 conflicts), indicating that the annotation error rate in these databases is ~17%.

## Background

Next-generation sequencing of markers such as the 16S ribosomal RNA (rRNA) gene and fungal internal transcribed spacer (ITS) region has revolutionized the study of microbial communities in environments ranging from the human body (Cho and Blaser, 2012; Pflughoeft and Versalovic, 2012) to oceans (Moran, 2015) and soils (Hartmann *et al.*, 2014). Specialized 16S rRNA sequence databases providing taxonomy annotations include Greengenes (DeSantis *et al.*, 2006), LTP (Yarza *et al.*, 2008), RDP (Maidak *et al.*, 2001), and SILVA (Pruesse *et al.*, 2007). LTP is based on sequences of type strains and isolates classified on the basis of observed traits. The other databases are much larger, containing mostly environmental sequences. In the RDP database, taxonomy annotations of environmental sequences are predicted by the Naive Bayesian Classifier (NBC) (Wang *et al.*, 2007), while those in Greengenes and SILVA are annotated by a combination of database-specific computational prediction methods and manual curation based on predicted phylogenetic trees inferred from multiple sequence alignments (McDonald *et al.*, 2012; Yilmaz *et al.*, 2014).

### Microbial taxonomy and phylogeny

The goal of taxonomy has been described as classifying living organisms into a hierarchy of categories (taxa) that are useful (based on biologically informative traits) and monophyletic (consistent with the true phylogenetic tree) (Hennig, 1965), though this point of view is not universally accepted (Benton, 2000). With microbial markers such as 16S rRNA, most organisms are known only from environmental sequencing, and in these cases predictions of taxonomy must necessarily be made from sequence evidence alone. One approach to predicting taxonomy is to infer a tree from available sequences, on the assumption that the tree will be a reasonable approximation to the phylogenetic tree and will therefore correlate with taxonomy. While the question of whether taxonomy should be consistent with phylogeny is controversial, the curators of Greengenes, LTP and SILVA have stated that they consider consistency to be important: “Greengenes is a dedicated full-length 16S rRNA gene database that provides users with a curated taxonomy based on *de novo* tree inference” (McDonald *et al.*, 2012); “[LTP is based on] comparative sequence analysis of SSU rRNA [which] has been established as the gold standard for reconstructing phylogenetic relationships among prokaryotes for classification purposes” (Yarza *et al.*, 2008); “SILVA predominantly uses phylogenetic classification based on an SSU guide tree … discrepancies are resolved with the overall aim of making classification consistent with phylogeny” (Yilmaz *et al.*, 2014). By contrast, taxonomy annotations for environmental sequences in RDP are predictions made by an algorithm, the Naive Bayesian Classifier, which does not consider phylogeny.

### Phylogenetic trees, gene trees and horizontal transfer

The history of *vertical* (i.e., clonal or sexual) inheritance can be represented as a binary tree. Inheritance can also occur by *horizontal* transfer of DNA between cells, which has been inferred to occur at a vast range of evolutionary distances ranging from members of the same species to exchanges between different kingdoms (Jain *et al.*, 2002). If a trait is acquired by horizontal transfer, then a graph representing its inheritance is not a tree. If horizontal transfer is rare, then in principle a phylogenetic tree could be constructed by following only vertically inherited traits. Gene content is sometimes highly variable between sets of prokaryotic genomes with otherwise highly similar sequences, which has been interpreted to imply that the optional genes which may be absent are frequently horizontally transferred (Gogarten and Townsend, 2005). If traits used for classification are acquired by horizontal transfer, then it may be impossible to achieve consistency between taxonomy and phylogeny. It is widely believed that the 16S rRNA gene is unlikely to be horizontally transferred and is thus well-suited as a phylogenetic marker (Woese, 1987), though recent evidence suggests that transfer of 16S rRNA genes can occur when sequence identity is as low as ~80% (Kitahara and Miyazaki, 2013). The evolutionary history of a single gene can be represented as a binary tree if its sequences are inherited as intact units, i.e. providing that rearrangement events such as cross-over recombinations do not occur. If rearrangement events and horizontal transfers are rare for the 16S rRNA gene, as is widely believed to be the case, then its true gene tree is likely to be a good approximation to the true phylogenetic tree based on vertically inherited traits, assuming that the latter tree can be meaningfully defined. However, even if there is good reason to believe that the true gene tree would be an accurate guide to taxonomy, it is very challenging to estimate the gene tree from multiple sequence alignments that span vast evolutionary distances, and it is an open question whether estimated gene trees enable accurate taxonomy prediction in practice.

### Taxonomy prediction from a sequence-based tree

Consider a binary tree with authoritative taxonomies for a small subset of its leaves. The goal is to predict taxonomies for the remaining leaves which represent organisms known only from sequence, based on the assumption that taxonomy is consistent with the tree. The *authoritative tree* is the implied binary tree for the subset of sequences with authoritative taxonomies, i.e. classifications based on direct observations of characteristic traits. Note that the annotations of this subset tree are considered to be authoritative, but not necessarily the branching order. The *lowest common ancestor* (LCA) of a taxon is the lowest node above all its leaves. If the branching order is correct and the taxonomy is consistent with the tree, then the subtree under the LCA for a taxon is *pure*, i.e. contains no other taxa at the same rank (this is the definition of consistency). Purity is closely related to monophyly; here I am making a distinction between consistency with a predicted tree which may have branching order errors (purity) and the true, but almost always unknown, phylogeny (monophyly). The *lowest pure taxon* (*LPT*) for a node is the taxon at lowest rank for which all the leaves in its subtree belong to the same taxon. For example, the LPT for node *n* is family *F* if, and only if, the leaves in the subtree under *n* belong to two or more genera in *F*. If *F* contains exactly one genus *G*, then *F* cannot be the LPT of any node because if *F* is pure then *G* will also be pure; *G* is lower, and the LPT would then be *G* rather than *F*. A *pure edge* connects two nodes having the same LPT. For example, consider the tree shown in Fig. 1. Sequences *s*, *g*_1_ and *g*_2_ are type strains with authoritative taxonomy; *w*, *x*, *y* and *z* are sequences obtained from environmental samples. The authoritative tree (black edges) connects *s*, *g*_1_ and *g*_2_ Genus is the lowest rank; *s* belongs to a singleton genus *S* (i.e., it is the only authoritatively named sequence for that genus); *g*_1_ and *g*_2_ belong to the same genus *G*. *S* and *G* belong to the same family, *F*. The LCA for *S* is *s* (a singleton leaf is always its own LCA), and the LCA for *G* is *g*. The LPTs of *g*_1_, *g*_2_ and *g* are *G*, the LPT of *s* is *S*, and the LPT of *f* is *F*. Pure edges are shown by continuous lines; impure edges are dashed lines. In general, if a tree is consistent with taxonomy, then all impure edges join an LCA node for some taxon *t* to a node which has an LPT at a higher rank than *t*, and conversely every LCA node has one impure edge which connects it to a higher node.

**Figure 1.**
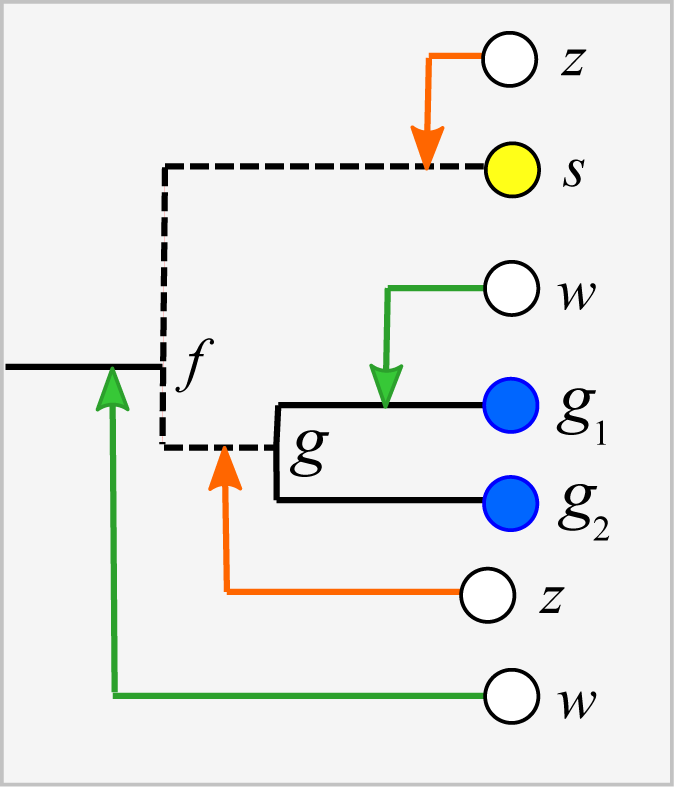
Example tree showing join nodes and impure edges. Leaves *s*, *g*_1_and *g*_2_ are sequences with authoritative taxonomies. Sequences *w*, *x*, *y* and *z* are environmental sequences. Sequence *s* belongs to genus *S*, which is a singleton (i.e., has only one ^authoritative sequence); sequences *g*_1_ and *g*_2_ belong to genus *G*. The LCA for *G* is *g*, the LCA for *S* is *s*.^ The inferred positions of *w*, *x*, *y* and *z* in the tree are indicated by arrows. An arrowhead is a join node, i.e., the new node created by adding a new sequence to the tree. Black edges are the authoritative tree, i.e., the binary tree for leaves with authoritative taxonomies. In the authoritative tree, dashed lines are impure edges; continuous lines are pure edges. If the join node is in a pure edge, the tree is sufficient to fully determine the taxonomy (green arrows); if the join node is in an impure edge, the tree is consistent with both a known or novel taxon (orange arrows).

#### The clade extension problem

Adding a new sequence (*Q*) of unknown taxonomy introduces a new node into the tree, its *join node*, indicated by an arrowhead in Fig. 1. Assume the tree is correct and the taxonomy is consistent with the tree. Then, if the join node is below the LCA for taxon *t*, *Q* certainly belongs to *t*. In Fig. 1, *y* certainly belongs to *G* because its join node is below *g*, the LCA of *G*. Also, *w* certainly belongs to a novel genus because it joins a family edge (which is above two or more genera by definition), and a new genus must therefore be introduced because it is impossible for the join node or any node above it to be a pure LCA of any known genus. The environmental sequence *x* joins the tree at the impure edge joining *s* and *f*, so it is above the LCA for *S*, but the subtree under the join node contains only *S*. The tree therefore cannot resolve whether *x* belongs to *S* or to a novel genus because both possibilities are consistent with the available data. If *x* belongs to *S*, its join node will be the new LCA for *S*, expanding the clade by moving its LCA higher in the tree. Otherwise, *x* belongs to a novel singleton genus, call it *V*, and *x* will be the LCA of *V* if it is added to the tree. Thus, *s* is the only sequence which unambiguously belongs to *S*—if a new sequence *Q* differs from *s*, then the join node is above the LCA for *S* and *Q* could belong to a different genus. This ambiguity could be resolved by introducing a sequence identity threshold. For example, if *x* is 97% identical to *s*, then it might be reasonable to assign *x* to *S*, but identity thresholds are only approximate guides and this prediction is uncertain even if the tree is assumed to be correct. Thus, an environmental sequence can be assigned to a singleton taxon only by considering sequence similarity, which is unreliable even under the idealized assumptions that the tree is correct and consistent with taxonomy. This is significant in practice because roughly half of named genera currently have only a single sequence in authoritative databases. A similar problem arises in non-singleton taxa. The join node for *z* is above the LCA for *G*, but the subtree below the join node contains only *G* and *z* could therefore belong to *G* (which would expand the clade by moving the LCA for *G* up to the join node for *z*), or to a novel genus, in which case *z* would be a new singleton LCA. In general, if an environmental sequence *Q* joins the tree at an impure edge, then the tree is consistent with two incompatible predictions: (1) assigning *Q* to a new taxon, or (2) extending an existing clade to include *Q*.

#### Impure taxa

Trees inferred from large sequence sets are unreliable guides to phylogeny due to incorrect branching orders (Huelsenbeck *et al.*, 2000; Philippe *et al.*, 2011). Naive assumptions that the tree is correct and the taxonomy is consistent with the tree are violated by branching order errors and also by taxa which are found to overlap by sequence such as *Escherichia* and *Shigella* (Escobar-Páramo *et al.*, 2003), which may be regarded as errors in the taxonomy. A conflict between the tree and the taxonomy occurs when taxa at the same rank overlap (Fig. 2). An *overlapped leaf* is below LCAs for two or more taxa at the same rank. Conversely, a taxon *t* is consistent with the tree if, and only if, the subtree under the LCA for *t* is pure. In Fig. 2, *i* is the LCA for *G* and *j* is the LCA for *H*; neither is pure because *i* is above *h*_1_ and *j* is above all members of both *G* and *H*. If an environmental sequence *Q* joins the tree at an edge where taxa overlap, i.e. the edge is above leaves belonging to two or more taxa at a given rank but below the LCAs for those taxa, the prediction at that rank is ambiguous.

**Figure 2.**
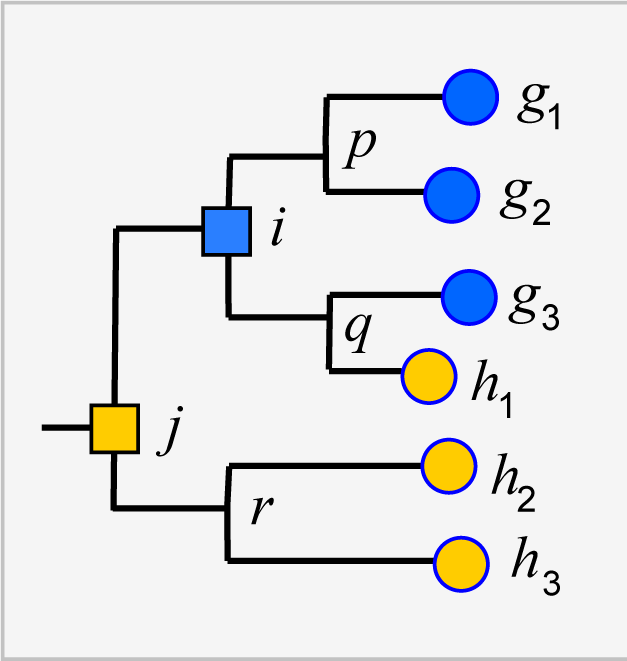
Example tree showing conflicts with taxonomy. Leaves *g*^1^, *g*_2_and *g*_3_ belong to genus *G*, *h*^1^, *h*_2_and *h*_3_ belong to a different genus *H*. The tree is not consistent with the taxonomy because *G* and *H* overlap. The LCA for *G* is *i*, which is above *h*_1_, and the LCA for *H* is *j*, which is above all members of both *G* and *H*. Overlaps are common in practice (Fig. 5).

For example, consider cases where the join node for *Q* is in *p-i*, *q-i, h*_1_-*q* or *g*_3_-*q*. Here, *Q* could plausibly belong to *G*, *H* or a novel genus, depending on the underlying explanation of the tree conflict. If *Q* joins the tree below a node for which taxon *t* is pure, then it is plausible that *Q* belongs to *t*, though this *purity heuristic* is not reliable because the branching order may be incorrect. For example, suppose *h*_1_ in Fig. 2 is an environmental sequence rather than a known strain, then the subtree under *i* is pure at genus rank and *h*_1_ would be incorrectly assigned to *G* by the purity heuristic.

### Taxonomic nomenclatures

At least three prokaryotic taxonomy nomenclatures are widely used. The Greengenes nomenclature is based on the NCBI Taxonomy database (Sayers *et al.*, 2012), RDP uses Bergey’s Manual (Bergey, 2001), while LPT and SILVA are based on LSPN (Parte, 2014). These nomenclatures differ mainly in revisions to resolve conflicts with sequence-based phylogenies and add new candidate groups identified in environmental sequences. For example, *Escherichia* and *Shigella* overlap as noted above, which Greengenes resolves by leaving their sequences unclassified at ranks below *Enterobacteriaceae.* SILVA, RDP and LTP deal with this issue differently by retaining well-established species names such as *Escherichia coli* and introducing a combined genus named *Escherichia-Shigella*. An example of an environmental candidate group is JS1 (Webster *et al.*, 2004), which has been adopted in the SILVA nomenclature but not in RDP or Greengenes at the time of writing.

## Methods

### Database versions

Unless otherwise stated, I used these databases: Greengenes v13.5, LTP v128, RDP downloaded on 15th January 2017, SILVA v128, the RDP 16S rRNA training set v16, and the BLAST 16S rRNA reference database (Sayers *et al.*, 2012) downloaded 17th January 2017 (BLAST16S).

### Common nomenclature

For a given pair of databases, I identified their *common nomenclature* as the set of taxon names appearing in both databases.

### Nomenclature hierarchy conflicts

Most well-established taxa are found in all nomenclatures, and in these cases a biologist would presumably expect a taxon name to be placed within the same group at a higher rank. To investigate this for a given pair of databases, I identified cases where a taxon name and its parent name were both in the common nomenclature in at least one of the databases, but the databases nevertheless disagreed on the parent name.

### Conflicting annotations of the same sequence

For a given pair of databases, I identified sequences that were present in both databases and were annotated as belonging to different taxa where at least one of the taxon names was in the common nomenclature.

### Conflicts between the LTP taxonomy and the LTP guide tree

For each taxon in the LTP nomenclature, I identified its LCA in the LTP tree (i.e., the lowest node above all members of the taxon), and then the overlapped taxa and leaves at each rank.

### Blinded testing of taxonomy predictions

If a sequence that was previously known only from environmental samples is found in a newly-sequenced isolate strain, it can provide a blinded test of taxonomy prediction analogous to CASP (Moult, 2005), where protein structures predicted from sequence are compared to newly-solved structures determined by sequence-independent methods such as X-ray crystallography. In principle, predictions for environmental sequences in an older version of Greengenes or SILVA could be assessed by finding identical sequences of isolates deposited more recently, but this approach is stymied by a lack of data specifying which sequences were used as references and the deposition dates of new isolate sequences. With RDP, it is straightforward because some of the older references (called training sets) for the RDP Naive Bayesian Classifier are deposited in public archives, and new sequences are readily identifiable as those not found in an older training set. If a training set is considered to be an authoritative reference, this enables predictions for new, authoritatively classified sequences to be assessed using an old training set as a reference, which is effectively equivalent to a blinded test. I implemented this test using versions 9 and 16 of the RDP 16S rRNA training sets, which were the oldest and newest, respectively, available for download at the time this work was performed. Version 9 contains 10,049 sequences; version 16 has 13,212 and therefore contains more than three thousand new sequences. I assessed predictions by NBC on these new sequences using v9 as a reference and the v16 annotations as the truth standard.

### Prediction performance metrics for the blinded test

I used the following metrics (Edgar, 2014, 2018) to measure prediction accuracy using a reference database (*A*) with a query set (*S*) containing sequences with known taxonomy. An *over-classification error* is a false positive where the taxon name of the query sequence is not present in the reference database. A *misclassification error* is a false positive where the taxon is known but the wrong name is predicted. An *under-classification error* is a false negative; i.e., case where no name is predicted but the taxon is known (present in *A*). Let *N* be the number of sequences in *S*, *K* be the number of query sequences with known names, i.e. names which are present in the reference database *A*, and *L* = *N* – *K* be the number of novel test sequences, i.e. sequences in *S* with names that are not present in *A*. Let *TP* be the number of names which are correctly predicted, *MC* the number of misclassification errors, *OC* the number of over-classification errors, and *UC* the number under-classification errors. The rate for each type of error is defined as the number of errors divided by the number of opportunities to make that error: *OCR = OC*/*L* (over-classification rate), *UCR* = *UC*/*K* (under-classification rate) and *MCR* = *MC/K* (misclassification rate). The true positive rate is *TPR* = *TP*/*K*, i.e. the number of correct names divided by the number of opportunities to correctly predict a name. *Accuracy* is calculated as *Acc* = *TP*/(*K* + *OC*), i.e. the number of correct predictions divided by the number of predictions for which correctness can be determined. Metrics are calculated separately for each rank.

### Conflicts between the RDP taxonomy and SSU guide trees

Greengenes and SILVA use sequence-based trees to guide predictions of taxonomy for environmental sequences. Conflicts between a phylogenetic tree and a taxonomic hierarchy arise when taxa at the same rank overlap in the tree (see Background). I used RDP training set v16 as a reference to assess the Greengenes guide tree from 2011 (http://greengenes.lbl.gov/Download/Trees/16S_all_gg_2011_1.tree.gz) and the SILVA guide tree from release 108 (dated Aug. 2011). This assesses older guide trees of approximately the same age by comparison with each other and with a more recent authoritative reference. As with the RDP blinded test, using an older version of the tree ensures that some newer isolate sequences are blinded, i.e., have authoritative taxonomies in the reference but predicted taxonomies in the trees. Using an independent reference minimizes bias due to manual adjustment of the trees to achieve consistency with reference sequences. I identified the *common subset* of sequences, i.e. sequences found in all three databases: SILVA, Greengenes and the training set. For each guide tree, I extracted the binary tree for the common subset of sequences (Fig. 3). The leaves in this subset tree were assigned taxonomy annotations according to the RDP training set while preserving the branching order specified by the guide tree. For each taxon in the RDP nomenclature, I identified its LCA in this subset tree (i.e., the lowest node above all members of the taxon), and then the overlapped taxa and leaves at each rank.

**Figure 3.**
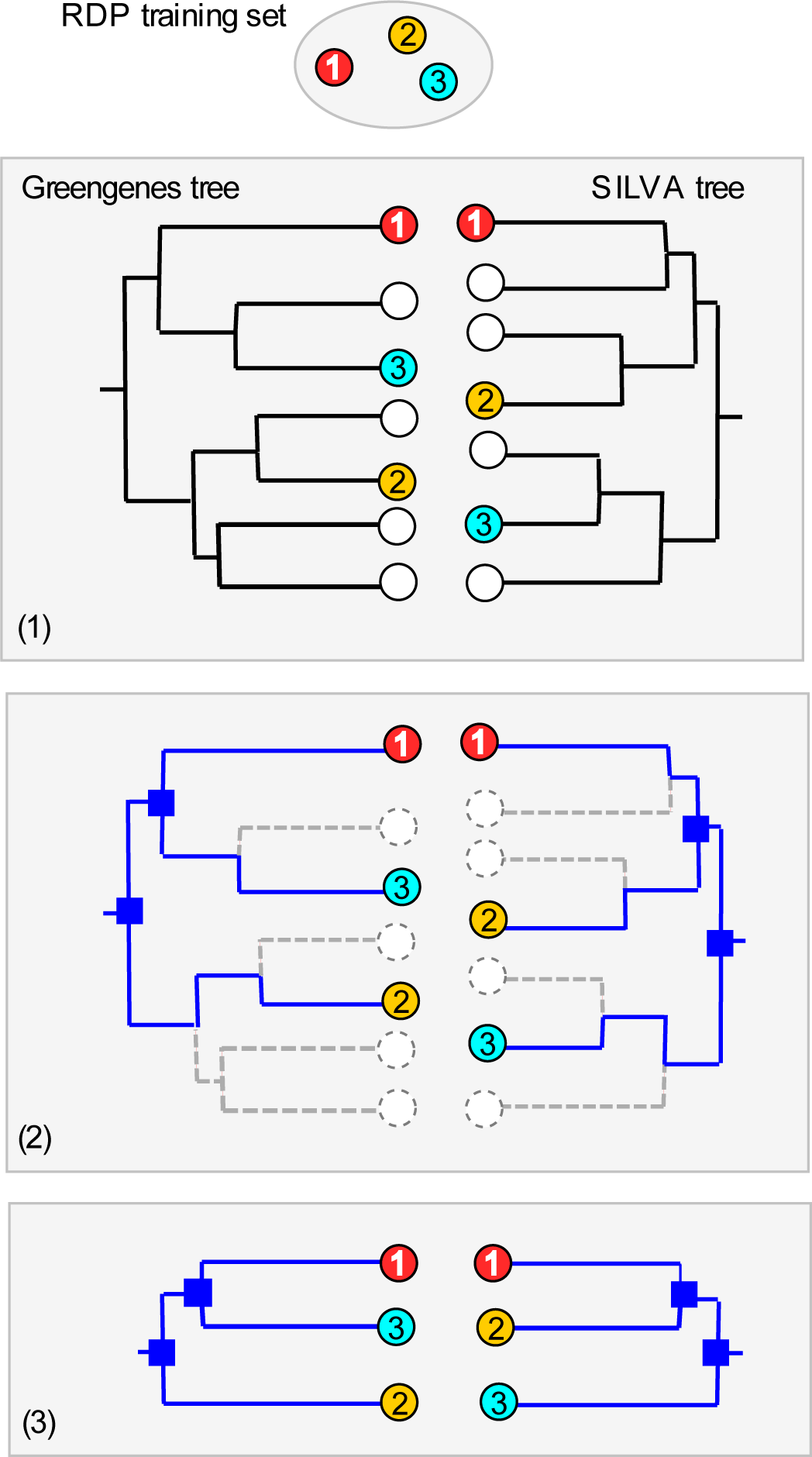
Concurrence of Greengenes and SILVA trees with RDP taxonomy. (1) Identical sequences (the common subset) are identified in all three databases (Greengenes, SILVA and the RDP training set). (2) Nodes and edges (shown in blue) specifying the branching order of the common subset are identified in each tree. (3) The binary trees implied by those nodes and edges are extracted and compared to each other and with the taxonomy annotations according to RDP. In this example, the branching order is different. If the three leaves belong to the same taxon, there is no conflict, otherwise one or both trees have a conflict with the RDP taxonomy.

### Consistency between SSU guide trees

A taxon that is impure according to an SSU guide tree implies one of the following: (1) the taxon is monophyletic and there are branching order errors in the tree, (2) the taxon is polyphyletic and the branching order correctly indicates the overlap, or (3) the taxon is in fact polyphyletic, but overlaps indicated by the branching order are different from those in the true phylogenetic tree. If case (2) is common, then subtrees of two independently-constructed guide trees should often be consistent with each other despite conflicts with taxonomy. To investigate whether this is the case in practice, I sought a suitable strategy for comparing two guide trees with taxonomy annotations for the same set of sequences. Published metrics for tree comparison include *Robinson-Foulds* (RF) (Robinson and Foulds, 1981) and *Branch Score Distance* (BSD) (Kuhner and Felsenstein, 1994), but these are not suitable here as the connection between their numerical values and taxonomic consistency is unclear, noting that branching errors which preserve purity are less important than those which cause monophyletic taxa to become impure in the predicted tree. It is more informative to measure the degree to which the trees agree on pure and overlapping taxa. For example, if trees *X* and *Y* agree that taxa *s* and *t* overlap, this suggests that both trees may be correctly implying that *s* and *t* overlap in the true phylogenetic tree, while if *s* and *t* are pure in *X* but overlap in *Y*, this suggests that the branching order of *X* is better than *Y*. I therefore defined the *Taxonomy Concurrence Score* (TCS) of trees *X* and *Y* as follows. Let LCA*X*(*t*) be the LCA of taxon *t* in tree *X* and let *SX*(*n*) be the set of taxon names at a given rank found in the subtree under node *n*. Then, *t* is pure in *X* if, and only if, *t* is the only taxon in *SX*(LCA*X*(*t*)). If *t* is impure, then there are two or more taxa in this set. Trees *X* and *Y* concur with respect to *t* if the same set of taxon names is found under the LCA for *t* in both trees, i.e. if *SX*(LCA*X*(*t*)) = *SY*(LCA*Y*(*t*)). Thus, the trees concur if *t* is pure in both trees, or the same set of overlapping taxon names is found under the LCA of *t*. An alternative design would be to require that the same set of sequences is found, but this would be excessively restrictive as it would treat a single misplaced sequence as equivalent to many disagreements. By considering names rather than sequences, the trees concur if they both imply that, say, *s* and *t* overlap with each other even if they place some sequences of *s* and *t* differently. TCS at a given rank is calculated as the fraction of non-singleton taxa where the trees concur. Singleton taxa are excluded because they are pure in all possible trees and are therefore uninformative.

### High-identity outliers

The *lowest common rank* (LCR) (Edgar, 2018) is the lowest rank shared by a pair of sequences. Note that this is the name of a rank, not a taxon name at that rank. For example, if two sequences belong to different genera in the same family, then their LCR is family. If a pair of sequences has an anomalously high identity considering its LCR, this indicates a possible classification error or annotation error, though it could also be explained by unusually high sequence conservation or by horizontal transfer. For example, if two 16S rRNA sequences are 99% identical and are annotated as belonging to different orders in the same class, this is likely to be a problem in the annotations.

## Results

### Annotation errors in the full RDP database

Annotations in the full RDP database are generated by the Naive Bayesian Classifier (NBC). Performance metrics for NBC predictions on new sequences in RDP training set v16 using v9 as a reference are shown in Table 1. The top-hit identity distribution (Edgar, 2018) of the blinded test is shown in Fig. 4, together with the distribution of the full RDP database against training set v16 for comparison. The full distribution is strongly peaked at 99% identity, while the blinded distribution has a higher frequency of low-identity sequences. The blinded test is therefore more challenging than annotating the full database, and the average genus accuracy of the RDP annotations is therefore probably higher than the ~85% observed in the blinded test. Given these results, a rough estimate is that ~10% sequences in RDP have incorrect genus annotations.

**Table 1.** Accuracy metrics for blinded RDP prediction test. See main text for definition of metrics.

| *Rank* | *TPR* | *MCR* | *OCR* | *UCR* | *Acc* |
| --- | --- | --- | --- | --- | --- |
| Phylum | 99.5% | 0.1% | 41.1% | 0.1% | 98.8% |
| Class | 99.1% | 0.4% | 90.2% | 0.4% | 83.5% |
| Order | 97.8% | 0.7% | 84.2% | 0.7% | 92.9% |
| Family | 96.9% | 0.7% | 71.7% | 0.7% | 92.8% |
| Genus | 91.8% | 2.6% | 41.9% | 2.6% | 84.8% |

**Figure 4.**
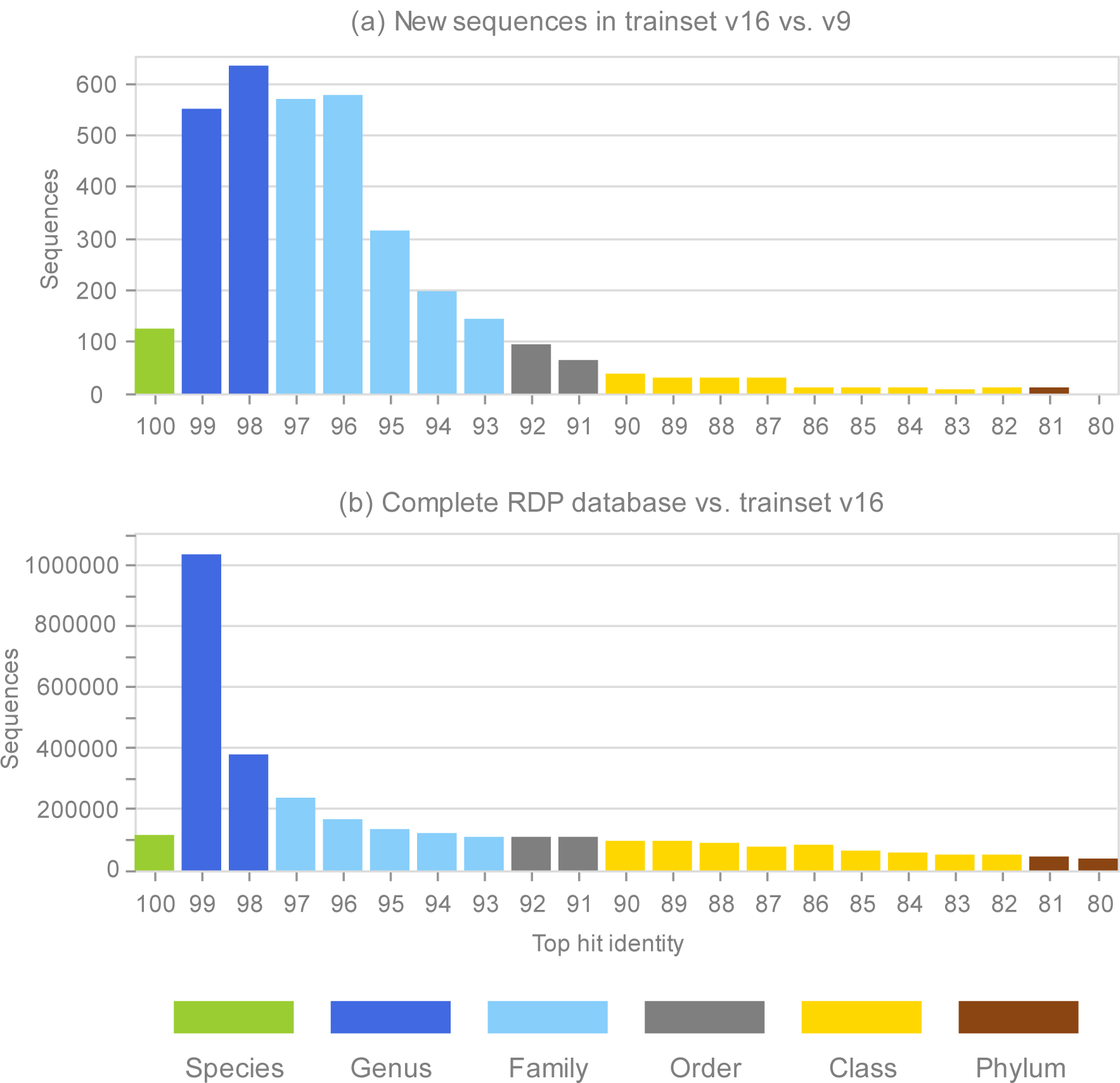
Top-hit identity distribution for the blinded RDP prediction test compared with the full RDP database. Panel (a) shows the top-hit identity distribution (THID) for new sequences in RDP trainset v16 compared to v9 as a reference. Panel (b) shows the THID for the full RDP database against trainset v16 as a reference. Histogram bars are colored according to the most probable lowest common rank (Edgar, 2018).

### Nomenclature hierarchy disagreements between Greengenes and SILVA

I found 46 taxon names placed into different parent taxa by Greengenes v13.5 and SILVA v128, considering only names present in both databases. For example, family *Brevibacteriaceae* is in order *Actinomycetales* according to Greengenes (e.g. accession 1109545), but this family is in order *Micrococcales* according to SILVA (JAJB01000072.80. 1512). Order *Micrococcales* is also found in Greengenes (825086), so there is an unambiguous disagreement between the nomenclature hierarchies. A complete list is given in Supp. Table 1.

### Annotation conflicts between Greengenes and SILVA

I found 784,242 identical sequences in SILVA and Greengenes, of which 732,048 (93%) had taxonomy annotations from the common nomenclature in one or both databases. Of the sequences with common nomenclature annotations, I found that 249,490 (34%) disagreed (Table 2). 59,637 (24%) of these conflicts were due to a rank which was blank (unspecified) in one database, which implies a false positive by one database or a false negative by the other. An example of a conflict between a pair of names in the common nomenclature is Greengenes 4366627 which has the same sequence as SILVA LOSM01000005.1106908.1108461. Greengenes assigns this sequence to family *Pseudoalteromonadaceae* while SILVA assigns it to family *Vibrionaceae*.

**Table 2.** Conflicts between Greengenes and SILVA annotations of identical sequences. Columns are: *Rank*, the highest rank where there was a disagreement between names in the common nomenclature; *Conflicts*, the number of disagreements (and as a percentage of annotations in the common nomenclature); *Blank*, the number of conflicts where the rank was not named in one of the databases; *Hierarchy*, conflicts where the databases disagreed on the parent name in cases where both the rank and its parent are in the common nomenclature in both databases.

| *Rank* | *Conflicts* | *Blank* | *Hierarchy* |
| --- | --- | --- | --- |
| Phylum | 7,804 (1.1%) | 92 | 0 |
| Class | 23,082 (3.2%) | 489 | 0 |
| Order | 110,308 (15.1%) | 2,777 | 4 |
| Family | 45,712 (6.2%) | 20,093 | 0 |
| Genus | 62,584 (8.5%) | 36,186 | 203 |
| Total | 249,490 (34.1%) | 59,637 | 207 |

*Pseudoalteromonadaceae* is found in SILVA (FJ889568.1.1501) and Greengenes contains *Vibrionaceae* (3059893). A small fraction of the conflicts (440, or 0.2%) reflected nomenclature hierarchy disagreements (see previous section); the rest were conflicts in taxon names for which both databases agreed on the parent group. Conflicts were found at all ranks: 7,804 at phylum, 23,082 class, 110,308 order, 45,712 family and 62,584 genus, counting only the highest conflicted rank. An example of a phylum conflict in a sequence classified to genus level by both databases is Greengenes 851746 which is assigned to genus *Streptococcus* in phylum *Firmicutes* and has the same sequence as SILVA JZIA01000003.3571786.3573313, which is assigned to genus *Xanthomonas* in phylum *Proteobacteria*. Assuming that a given named taxon is required to have the same definition (e.g., the same characteristic traits) in all nomenclatures where it appears, then all conflicts are necessarily due to annotation errors in one or both databases. The lower bound on the sum of the number of errors in both databases is obtained when all conflicts are due to an error in one database but not the other. This follows because cases where both databases make the same error (so there is no disagreement, but the annotations are nevertheless wrong), or a given sequence has errors in both databases, can only add to this sum. Thus, a lower bound for the sum of the annotation error rates of SILVA and Greengenes is (number of identical sequences with common nomenclature conflicts) / (number of identical sequences annotated by the common nomenclature) = 249,490/732,048 = 34%.

### Conflicts between the LTP taxonomy and the LTP guide tree

Result for genus and family are shown in Table 3; higher ranks are not annotated in LTP. At genus rank, roughly one-third of non-singleton genera overlap (430/1,328 = 32%) and at family rank almost one-half overlap (155/326 = 48%). A large majority of leaves (~300/12,953 = 98%) are overlapped at both family and genus rank, i.e. are in the subtrees under LCAs for two or more taxa at the same rank.

**Table 3.** Conflicts between the tree and taxonomy annotations in LTP. LTP does not annotate ranks above genus. Columns are: *Singletons*, the number of singleton taxa; *Non-singletons*, the number of non-singleton taxa; *Pure taxa*, the number of non-singleton taxa which are pure; *Impure taxa*, the number of non-singleton taxa which are impure, *Pure leaves*, the number of pure leaves, i.e. the number of leaves found under the LCA for exactly one taxon; *Impure leaves*, the number of leaves which are not pure.

|  | *Genus* | *Family* |
| --- | --- | --- |
| *Singletons* | 1,206 | 79 |
| *Non-singletons* | 1,328 | 326 |
| *Pure taxa* | 898 (68%) | 171 (52%) |
| *Impure taxa* | 430 (32%) | 155 (48%) |
| *Pure leaves* | 273 | 295 |
| *Impure leaves* | 12,680 | 12,658 |

### Common subset of SILVA, Greengenes and the RDP training set

Using the method sketched in Fig. 3, I found 7,216 identical sequences in SILVA, Greengenes and the RDP training set. Extracting the subtrees for these sequences from the SILVA and Greengenes guide trees gave subset trees with 7,216 leaves with taxonomy annotations according to the RDP training set.

### Concurrence of Greengenes and SILVA guide trees

Results are shown in Table 4. This shows the concurrence between the subset of the Greengenes and SILVA guide trees for sequences found in Greengenes, SILVA and the RDP training set, using taxonomy annotations from the training set. These subset trees are directly comparable because they contain the same set of sequences with taxonomy annotations from an independent database. The results show comparable rates of disagreement with the training set taxonomy. For example, 58 genera are pure according to the Greengenes tree but not SILVA, while 74 genera are pure according to SILVA but not Greengenes. If one tree (call it *X*) was substantially better than the other (*Y*), then there would be more taxa that are impure in *Y* but not in *X* than *vice versa*, but this is not the case here, which indicates that the Greengenes and SILVA trees have comparable accuracy, as might be expected. There are 74 genera which both trees agree are impure and have the same set of overlapping names. These cases could be explained by taxa which are truly polyphyletic, or by branching order errors which are reproduced in both trees. Reproduced errors may occur by chance, noting that noise which is unbiased when measured by branching order will have a strong tendency to merge neighboring taxa, and will therefore be strongly biased when measured by taxonomy. Reproduced errors may also occur due to systematic effects such as long branch attraction, which is known to occur in a wide range of tree inference algorithms including neighbor-joining, parsimony, maximum likelihood, and Bayesian methods (Bergsten, 2005).

**Table 4.** Concurrence between Greengenes, SILVA and RDP training set. Concurrence for identical sequences found in all three databases determined by the method shown in Fig. 3. Columns are: *Pure both*, the number of non-singleton taxa which are pure in the subset trees for both Greengenes and SILVA; *Pure GG only*, the number which are pure only in the Greengenes subset tree; *Pure SILVA only*, similarly for SILVA; *Impure, same set*, the number of taxa where the trees agreed on the set of overlapping names; *Impure, diff. set*, the number where the trees disagreed on the set of overlapping names; *TCS*, taxonomy concurrence score, i.e. the fraction of taxa where the trees concur that the taxon was pure or had the same set of overlapping names under its LCA.

| *Rank* | *Pure both* | *Pure*<br>*GG only* | *Pure*<br>*SILVA only* | *Impure,*<br>*same set* | *Impure,*<br>*diff. set* | *TCS* |
| --- | --- | --- | --- | --- | --- | --- |
| Phylum | 37 (78.7%) | 0 (0.0%) | 3 (6.4%) | 3 (6.4%) | 4 (8.5%) | 40 (85.1%) |
| Class | 65 (79.3%) | 2 (2.4%) | 2 (2.4%) | 6 (7.3%) | 7 (8.5%) | 71 (86.6%) |
| Order | 89 (65.0%) | 7 (5.1%) | 3 (2.2%) | 12 (8.8%) | 26 (19.0%) | 101 (73.7%) |
| Family | 198 (61.3%) | 20 (6.2%) | 13 (4.0%) | 31 (9.6%) | 61 (18.9%) | 229 (70.9%) |
| Genus | 1449 (82.5%) | 58 (3.3%) | 74 (4.2%) | 74 (4.2%) | 101 (5.8%) | 1523 (86.7%) |

### Rhodobacter subtrees

The subtrees for genus *Rhodobacter* in Greengenes and SILVA are shown in Fig. 5. After manually reviewing many taxa, I selected *Rhodobacter* as a representative example with different branching orders. As is typically the case, groups of highly similar sequences are concurrent between the trees. For example, *Rhodobacter* sequences X54853, AM398152 and AM421024 have pair-wise identities >99% and are placed together in both trees. When sequence similarity is lower, there is often radical disagreement in the branching order. For example, the two *Heamobacter* sequences AF45106 and DQ342315 are placed inside the *Gemmobacter* subtree by Greengenes, while both *Haemobacter* and *Gemmobacter* are pure according to SILVA. With the LTP tree, the subtree under its *Rhodobacter* LCA node contains 390 sequences from 122 genera as shown in Supp. Fig. 1.

**Figure 5.**
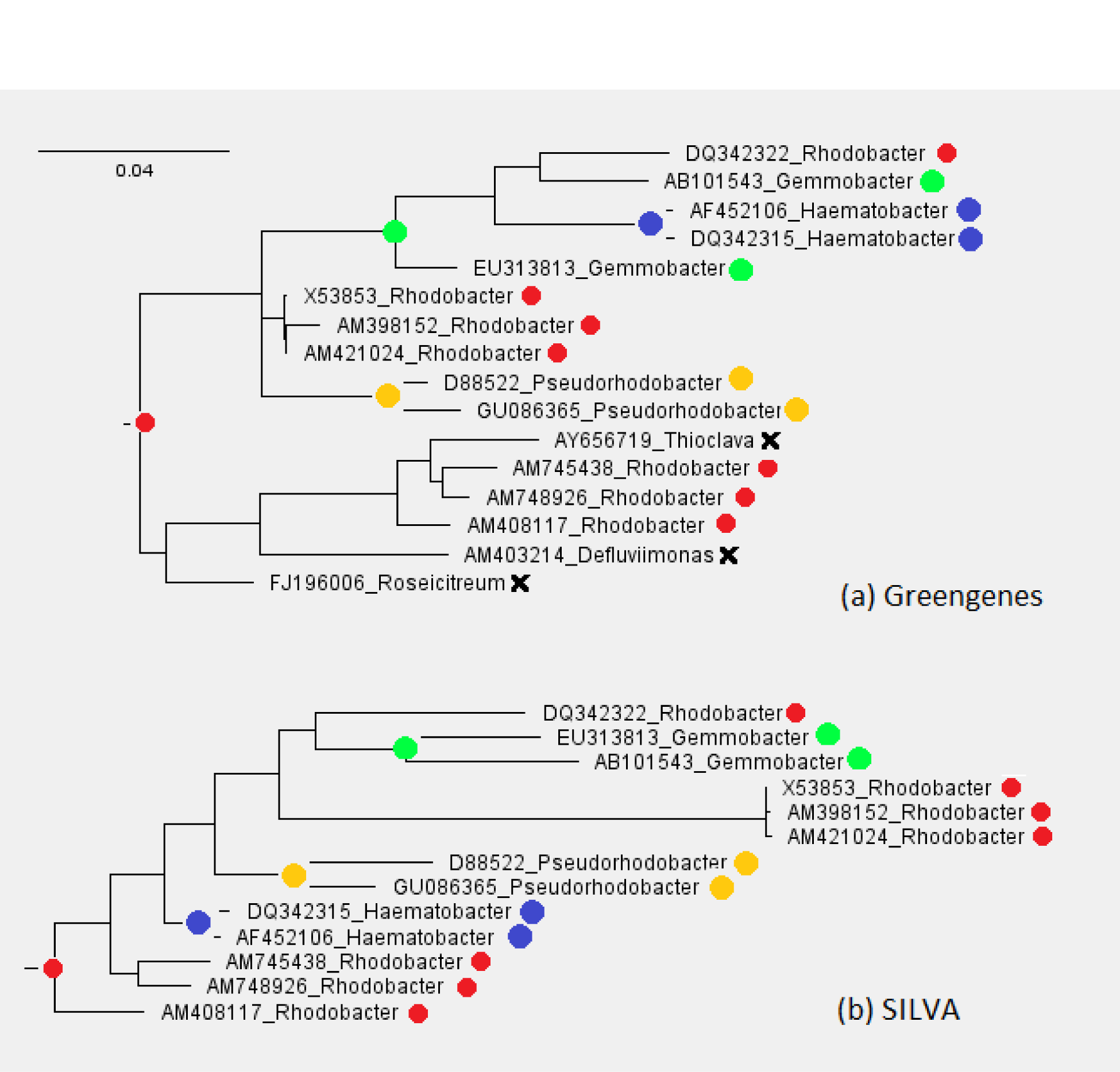
Subtrees for *Rhodobacter* in Greengenes and SILVA. Leaves are labeled with their sequence identifiers and genus names according to the RDP training set. Colored dots indicate genera found in both subtrees; black crosses indicate genera found in the Greengenes subtree only.

### High-identity outliers

The outliers with highest pair-wise identity at each LCR for BLAST16S and the RDP training set are given in Tables 5 and 6 respectively. The ten pairs with highest identity at each rank for all these databases and also LTP are given in Supp. Table 2. As an example, sequences gi|645320505 and gi|265678369 in BLAST16S are 100% identical and have LCR=phylum, which almost certainly implies an annotation error.

**Table 5.** Outliers with high pair-wise identity in the BLAST16S database. Columns are: *%id*, identity; *LCR*, lowest common rank; *Accessions*, database identifiers; *Highest Distinct Taxa*, names at the highest rank where the sequences are assigned to different taxa; *Species*, species names according to the database.

| *%Id* | *LCR* | *Accessions* | *Highest Distinct Taxa* | *Species* |
| --- | --- | --- | --- | --- |
| 100.0 | Phylum | gi\|645320505<br>gi\|265678369 | c:Betaproteobacteria<br>c:Gammaproteobacteria | *Neisseria flava*<br>*Moraxella caviae* |
| 97.1 | Phylum | gi\|559795246<br>gi\|645321781 | c:Bacilli<br>c:Clostridia | *Lactobacillus rogosae*<br>*Lachnospira pectinoschiza* |
| 99.9 | Class | gi\|558508609<br>gi\|631251240 | o:Pseudonocardiales<br>o:Streptomycetales | *Prauserella marina*<br>*Streptomyces parvus* |
| 99.6 | Class | gi\|645319747<br>gi\|343198728 | o:Pseudomonadales<br>o:Oceanospirillales | *Pseudomonas halophila*<br>*Halovibrio denitrificans* |
| 100.0 | Order | gi\|636559468<br>gi\|219846253 | f:Bacillaceae<br>f:Sporolactobacillaceae | *Bacillus racemilacticus*<br>*Sporolactobacillus laevolacticus* |
| 100.0 | Order | gi\|645320480<br>gi\|343200250 | f:Rhodobiaceae<br>f:Bradyrhizobiaceae | *Afifella marina*<br>*Rhodopseudomonas julia* |
| 100.0 | Family | gi\|636560543<br>gi\|631251787 | g:Obesumbacterium<br>g:Hafnia | *Obesumbacterium proteus*<br>*Hafnia alvei* |
| 100.0 | Family | gi\|1212229262<br>gi\|1137647790 | g:Feifantangia<br>g:Xanthomarina | *Feifantangia zhejiangensis*<br>*Xanthomarina gelatinilytica* |
| 100.0 | Genus | gi\|343198889<br>gi\|343201672 | s:aurantiaca<br>s:erecta | *Stigmatella aurantiaca*<br>*Stigmatella erecta* |
| 100.0 | Genus | gi\|219857540<br>gi\|219846097 | s:florum<br>s:grammopterae | *Mesoplasma florum*<br>*Mesoplasma grammopterae* |

**Table 6.** Outliers with high pair-wise identity in the RDP training set. Columns as for Table 5.

| *%Id* | *LCR* | *Accessions* | *Highest Distinct Taxa* | *Genus* |
| --- | --- | --- | --- | --- |
| 98.1 | Phylum | AB680539<br>Y10109 | c:Betaproteobacteria<br>c:Alphaproteobacteria | *Aquaspirillum*<br>*Magnetospirillum* |
| 99.9 | Class | Y10755<br>AB021399 | o:Xanthomonadales<br>o:Pseudomonadales | *Xanthomonas*<br>*Pseudomonas* |
| 99.6 | Class | AJ007800<br>AJ227812 | o:Caulobacterales<br>o:Sphingomonadales | *Asticcacaulis*<br>*Sphingomonas* |
| 99.8 | Order | AY345990<br>FR733691 | f:Entomoplasmataceae<br>f:Spiroplasmataceae | *Entomoplasma*<br>*Spiroplasma* |
| 99.1 | Order | AJ006086<br>EF114311 | f:Planococcaceae<br>f:Bacillaceae_1 | *Solibacillus*<br>*Bacillus* |
| 100.0 | Family | GU256437<br>KC109787 | g:Nitrospirillum<br>g:Azospirillum | *Nitrospirillum*<br>*Azospirillum* |
| 99.9 | Family | FR733721<br>X79289 | g:Nocardia<br>g:Rhodococcus | *Nocardia*<br>*Rhodococcus* |

## Discussion

The results presented show that guide trees inferred from 16S rRNA sequences have pervasive conflicts with taxonomy. Guide trees in different databases conflict with each other and with taxonomy at comparable rates, which strongly suggests that many or most of these conflicts are due to branching order errors in the predicted trees compared with the true gene tree. A lower bound of 34% was obtained for the sum of annotation error rates in Greengenes and SILVA. I believe that this bound is robust because it is obtained from names in their common nomenclature, so any disagreement is necessarily due to an error in one or both annotations, and a large majority (>90%) are have common nomenclature annotations. The number of conflicts between the guide trees and an independent reference (the RDP training set) is similar, and neither tree contains substantially more pure taxa according to the RDP training set. These results imply that the annotation error rate of both databases is similar, as might be expected. If the error rates are similar and the sum is close to its lower bound of 34%, then the annotation error rates of Greengenes and SILVA are both approximately 17%. If one database is somewhat better (say, its error rate is 15%) then the error rate of the other must be correspondingly higher (19%) so that the sum remains at least 34%. It seems very unlikely that either database could have an error rate as low as 10% as this would imply that the other has an error rate of at least 24%, and a difference of this magnitude should be noticeable in the numbers of pure taxa. Noting that the sum may be somewhat higher than its lower bound due to systematic errors reproduced in both databases, a reasonable rough estimate is that one in five sequences in Greengenes and SILVA have incorrect taxonomy annotations. The blinded test suggests that the annotation rate of RDP is ~10% and is therefore likely to be lower than Greengenes or SILVA, though the uncertainties in these estimates are too great to support a firm conclusion that annotations in any one of these databases are better or worse than another. However, while keeping this caveat in mind, it is striking that a pessimistic estimate for the RDP database of 15% on the blinded test is less than a conservative estimate for SILVA and Greengenes of a 17% error rate from disagreements between annotations. This suggests that the accuracy of fully-automated annotations generated by the RDP Naive Bayesian Classifier, which does not explicitly consider phylogeny, is quite likely to be better than the accuracy of annotations in SILVA and Greengenes, which were generated by a combination of automated and manual methods guided by alignments and trees. Predictions generated by the RDP Classifier have several advantages: they are based on a documented reference, can easily be reproduced, and report a bootstrap confidence value. By contrast, annotations in SILVA and Greengenes do not report confidence and are not reproducible because there is a manual curation step and references are not documented.

## References

Benton MJ. (2000). Stems, nodes, crown clades, and rank-free lists: Is Linnaeus dead? Biol Rev 75: 633–648.

Bergey DH. (2001). Bergey’s manual of systematic bacteriology.

Bergsten J. (2005). A review of long-branch attraction. Cladistics 21: 163–193.

Cho I, Blaser MJ. (2012). The human microbiome: at the interface of health and disease. Nat Rev Genet 13: 260–270.

DeSantis TZ, Hugenholtz P, Larsen N, Rojas M, Brodie EL, Keller K, et al. (2006). Greengenes, a chimera-checked 16S rRNA gene database and workbench compatible with ARB. Appl Environ Microbiol 72: 5069–72.

Edgar RC. (2018). Accuracy of taxonomy prediction for 16S rRNA and fungal ITS sequences. PeerJ. Edgar RC. (2014). Taxonomy benchmark tests, USEARCH manual v8.1. https://drive5.com/usearch/manual8.1.

Escobar-Páramo P, Giudicelli C, Parsot C, Denamur E. (2003). The evolutionary history of Shigella and enteroinvasive Escherichia coli revised. J Mol Evol 57: 140–148.

Gogarten JP, Townsend JP. (2005). Horizontal gene transfer, genome innovation and evolution. Nat Rev Microbiol 3: 679–687.

Hartmann M, Niklaus PA, Zimmermann S, Schmutz S, Kremer J, Abarenkov K, et al. (2014). Resistance and resilience of the forest soil microbiome to logging-associated compaction. ISME J 8: 226–44.

Hennig W. (1965). Phylogenetic Systematics. Annu Rev Entomol 10: 97–116.

Huelsenbeck JP, Rannala B, Masly JP. (2000). Accommodating phylogenetic uncertainty in evolutionary studies. Science (80-) 288: 2349–2350.

Jain R, Rivera MC, Moore JE, Lake JA. (2002). Horizontal Gene Transfer in Microbial Genome Evolution. Theor Popul Biol 61: 489–495.

Kitahara K, Miyazaki K. (2013). Revisiting bacterial phylogeny: Natural and experimental evidence for horizontal gene transfer of 16S rRNA. Mob Genet Elements 3: e24210.

Kuhner MK, Felsenstein J. (1994). A simulation comparison of phylogeny algorithms under equal and unequal evolutionary rates. Mol Biol Evol 11: 459–468.

Maidak BL, Cole JR, Lilburn TG, Parker CT, Saxman PR, Farris RJ, et al. (2001). The RDP-II (Ribosomal Database Project). Nucleic Acids Res 29: 173–4.

McDonald D, Price MN, Goodrich J, Nawrocki EP, DeSantis TZ, Probst A, et al. (2012). An improved Greengenes taxonomy with explicit ranks for ecological and evolutionary analyses of bacteria and archaea. ISME J 6: 610–618.

Moran MA. (2015). The global ocean microbiome. Science (80-) 347: aac8455.

Moult J. (2005). A decade of CASP: Progress, bottlenecks and prognosis in protein structure prediction. Curr Opin Struct Biol 15: 285–289.

Parte AC. (2014). LPSN - List of prokaryotic names with standing in nomenclature. Nucleic Acids Res 42. e-pub ahead of print, doi: 10.1093/nar/gkt1111.

Pflughoeft KJ, Versalovic J. (2012). Human microbiome in health and disease. Annu Rev Pathol 7: 99–122.

Philippe H, Brinkmann H, Lavrov D V., Littlewood DTJ, Manuel M, Wörheide G, et al. (2011). Resolving difficult phylogenetic questions: Why more sequences are not enough. PLoS Biol 9. e-pub ahead of print, doi: 10.1371/journal.pbio.1000602.

Pruesse E, Quast C, Knittel K, Fuchs BM, Ludwig W, Peplies J, et al. (2007). SILVA: A comprehensive online resource for quality checked and aligned ribosomal RNA sequence data compatible with ARB. Nucleic Acids Res 35: 7188–7196.

Robinson DF, Foulds LR. (1981). Comparison of phylogenetic trees. Math Biosci 53: 131–147.

Sayers EW, Barrett T, Benson DA, Bolton E, Bryant SH, Canese K, et al. (2012). Database resources of the National Center for Biotechnology Information. Nucleic Acids Res 40: D13–25.

Wang Q, Garrity GM, Tiedje JM, Cole JR. (2007). Naive Bayesian classifier for rapid assignment of rRNA sequences into the new bacterial taxonomy. Appl Environ Microbiol 73: 5261–7.

Webster G, Parkes RJ, Fry JC, Weightman AJ. (2004). Widespread occurrence of a novel division of bacteria identified by 16S rRNA gene sequences originally found in deep marine sediments. Appl Environ Microbiol 70: 5708–5713.

Woese C. (1987). Bacterial evolution background. Microbiology 51: 221–271.

Yarza P, Richter M, Peplies J, Euzeby J, Amann R, Schleifer KH, et al. (2008). The All-Species Living Tree project: A 16S rRNA-based phylogenetic tree of all sequenced type strains. Syst Appl Microbiol 31: 241–250.

Yilmaz P, Parfrey LW, Yarza P, Gerken J, Pruesse E, Quast C, et al. (2014). The SILVA and ‘all-species Living Tree Project (LTP)’ taxonomic frameworks. Nucleic Acids Res 42. e-pub ahead of print, doi: 10.1093/nar/gkt1209.

